# Hybrid Hyperalignment: A single high-dimensional model of shared information embedded in cortical patterns of response and functional connectivity

**DOI:** 10.1101/2020.11.25.398883

**Authors:** Erica L. Busch, Lukas Slipski, Ma Feilong, J. Swaroop Guntupalli, Matteo Visconti di Oleggio Castello, Jeremy F. Huckins, Samuel A. Nastase, M. Ida Gobbini, Tor D. Wager, James V. Haxby

**Affiliations:** Department of Psychology, Yale University, New Haven, CT, USA; Department of Psychological and Brain Sciences, Dartmouth College, Hanover, NH, USA; Vicarious AI, Union City, CA, USA; Helen Wills Neuroscience Institute, University of California, Berkeley; Princeton Neuroscience Institute, Princeton University, Princeton, NJ, USA; Department of Experimental, Diagnostic, and Specialty Medicine, Medical School, University of Bologna, Italy; Cognitive Science Program, Dartmouth College, Hanover, NH, USA

**Keywords:** fMRI, functional alignment, hyperalignment, naturalistic stimuli, functional connectivity.

## Abstract

Shared information content is represented across brains in idiosyncratic functional topographies. Hyperalignment addresses these idiosyncrasies by using neural responses to project individuals’ brain data into a common model space while maintaining the geometric relationships between distinct activity patterns. The dimensions of this common model can encode any kind of functional profiles shared across individuals, such as cortical response profiles collected during a common time-locked stimulus presentation (e.g. movie viewing) or functional connectivity profiles. Performing hyperalignment with either response-based or connectivity-based input data derives transformations to project individuals’ neural data from anatomical space into the common model such that functional information is optimally aligned across brains. Previously, only response or connectivity profiles were used in the derivation of these transformations. In this study, we used three separate data sets collected while participants watched feature films to derive transformations representing both response-based and connectivity-based information with a single algorithm. Our new method, hybrid hyperalignment, aligns response-based information as well as or better than response hyperalignment while simultaneously aligning connectivity-based information better than connectivity hyperalignment, all in one information space. These results suggest that a single common information space could encode both shared cortical response and functional connectivity profiles across individuals.

## 1 Introduction

Hyperalignment models shared information that is embedded in idiosyncratic cortical patterns across brains. The utility of modeling this shared information is that it makes possible much more accurate comparisons of functional activity across brains. Hyperalignment projects cortical pattern vectors into a common, high-dimensional information space (Haxby et al., 2020).Derivation of this common space can be based on either neural response profiles (e.g. data collected during tasks, such as movie viewing (Haxby et al., 2011) or functional connectivity profiles (Guntupalli et al., 2018). Common spaces based on each of these data types differentially improve between-subject alignment, with response-based common spaces better aligning held-out response data, and connectivity-based common spaces better aligning held-out connectivity data. However, it has remained unclear whether optimizations of both response hyperalignment and connectivity hyperalignment would converge on the same common information space.

While both response- and connectivity-based hyperalignment significantly improve the intersubject correlations (ISCs) of response profiles relative to anatomical alignment, response-based hyperalignment (RHA) results in slightly higher ISCs for response profiles than does connectivity-based hyperalignment (CHA) (Guntupalli et al., 2018). Similarly, RHA yields better alignment of cortical response patterns for two additional tests of between-subject alignment: between-subject multivariate pattern classification (bsMVPC) and ISC of representational geometry (Guntupalli et al., 2016, 2018). At the same time, CHA yields higher ISCs of dense connectivity profiles than RHA (Guntupalli et al., 2018). In other words, RHA outperforms CHA on response-based metrics of alignment whereas CHA outperforms RHA on connectivity-based metrics. The common information spaces derived from RHA and CHA are correlated yet different, suggesting that information contained in population response patterns and information contained in functional connectomes is fundamentally distinct. Alternatively, RHA and CHA may both be imperfect estimates of a single common information space that can accommodate both shared response information and shared connectivity information.

If the first hypothesis holds, and the common spaces derived by RHA and CHA each capitalize on distinct aspects of the same data, then two separate optimal common spaces exist. In this case, adding response information to connectivity-based hyperalignment would move the CHA common space toward the RHA optimum and away from the optimal CHA space, degrading ISC of connectivity profiles. Likewise, moving closer to the shared CHA space by adding connectivity information to response-based hyperalignment should degrade response-based benchmarks of between-subject alignment: ISC of response profiles and bsMVPC of response patterns.However, if the second hypothesis holds, both RHA and CHA are imperfect estimates of a single optimal shared-information space. In this case, deriving a common space based on combined response and connectivity data should maintain or improve ISCs of response and connectivity profiles as well as bsMVPC of response patterns.

We tested these two possibilities using fMRI data collected while participants watched one of three movies: *The Grand Budapest Hotel* (Visconti di Oleggio Castello, Chauhan, et al., 2020), *Raiders of the Lost Ark,* or *Whiplash*. We found that a single common model computed using both response and functional connectivity information aligned neural response and connectivity patterns across participants as well as or better than RHA or CHA alone, supporting the second hypothesis of a single, optimal shared-information space.

## 2 Materials and Methods

### 2.1 Participants

We used three separate data sets for our analyses. All participants gave written, informed consent, and all studies were approved by the Institutional Review Board of Dartmouth College. In data set one (Budapest), we scanned 21 participants (11 female, 27.29 years ± 2.35 SD) as they watched the second half of the film *The Grand Budapest Hotel* (Visconti di Oleggio Castello, Chauhan, et al., 2020). This dataset had 25 total participants, but we used a subset of 21 participants with headcases for this analysis. In data set two (Raiders), we scanned 23 participants (12 female, 27.26 years ± 2.40 SD) as they watched the second half of the film *Raiders of the Lost Ark*. In the third study (Whiplash), 29 participants (15 female, 18.30 years ±0.79 SD) watched part of the film *Whiplash*. In the Whiplash data set, the 29 participants with the least head motion, measured using average framewise displacement, were chosen from a set of 62 participants who viewed this video as part of another study.

### 2.2 Stimuli and Design

In each of these studies, participants viewed part of an audio-visual film in the MRI scanner. In the Budapest data set, participants watched the audio-visual film *The Grand Budapest Hotel*. They viewed the first portion of the movie outside of the scanner and the second portion (final 50.9 minutes) in the scanner as we collected fMRI data. This second portion of the film was broken into 5 separate runs, each approximately 10 minutes long, with a short break between each run (Visconti di Oleggio Castello, Chauhan, et al., 2020). In the Raiders data set, fMRI responses were measured while participants watched the second half of the film *Raiders of the Lost Ark* (approximately 57 minutes) over 4 runs, each roughly 15 minutes. Again, participants viewed the first half of the movie outside of the scanner just prior to the scanning session. In the Whiplash data set, participants watched a 29.5-minute edit of the film *Whiplash*. FMRI data was collected during all 29.5 minutes in a single run.

For each data set, the videos were projected using an LCD projector, which the participant could view on a mirror mounted on the head coil in the scanner. Audio was played using MRI-compatible in-ear headphones. Participants were simply instructed to pay attention and enjoy the movie.

### 2.3 MRI Data Acquisition and Preprocessing

All fMRI data were collected in the Dartmouth Brain Imaging Center with a 3T Siemens Magnetom Prisma MRI scanner (Siemens, Erlangen, Germany) with a 32-channel phased-array head coil with TR/TE = 1000/33 ms, flip angle = 59°, resolution = 2.5×2.5×2.5 mm isotropic voxels, matrix size = 96×96, FoV = 240×240 mm, with anterior-posterior phase encoding. For Budapest and Whiplash 52 axial slices were obtained. For Raiders 48 axial slices were obtained. Both volumes provided roughly full brain coverage with no gap between slices.

Anatomical data were acquired using a high-resolution 3-D magnetization-prepared rapid gradient echo sequence (MP-RAGE; 160 sagittal slices; TR/TE, 9.9/4.6 ms; flip angle, 8°; voxel size, 1×1×1 mm). Data acquisition and conversion to BIDS was performed using the ReproIn specification and tools (Visconti di Oleggio Castello, Dobson, et al., 2020) and organized into BIDS format with DataLad (Gorgolewski et al., 2016; Halchenko et al., 2017). Data was preprocessed using fMRIprep 20.0.3 (Esteban et al., 2018). The Budapest, Raiders, and Whiplash data sets had 3,052, 2,570, and 1,770 total TRs respectively.

### 2.4 Intersubject Alignment

Our analysis consisted of four types of intersubject alignment beginning with traditional anatomical alignment described in the previous section (and displayed in Fig. 1). Anatomical alignment (AA) nonlinearly registered each participant’s individual BOLD response data to FreeSurfer’s fsaverage7 cortical template based on sulcal curvature (Fischl, 2012). For computational efficiency, we then decimated this data to fsaverage5 by selecting the first 10,242, vertices per hemisphere. The AA data was then used to perform hyperalignment with three different algorithms. Response-based hyperalignment (RHA) mapped data from the fsaverage anatomical space to a common information space based on the movie-viewing response patterns across cortical vertices. Connectivity-based hyperalignment (CHA) mapped data from the fsaverage anatomical space to a separate common information space based on functional connectivity patterns derived from the movie response data. Finally, the novel hybrid hyperalignment (H2A) algorithm combined the input data used in both RHA and CHA and then calculated a third common information space based on both movie-viewing response patterns and the functional connectivity patterns derived therefrom. All hyperalignment was performed with python code utilizing the PyMVPA toolbox version 2.6.5 (Hanke et al., 2009).

**Figure 1:**
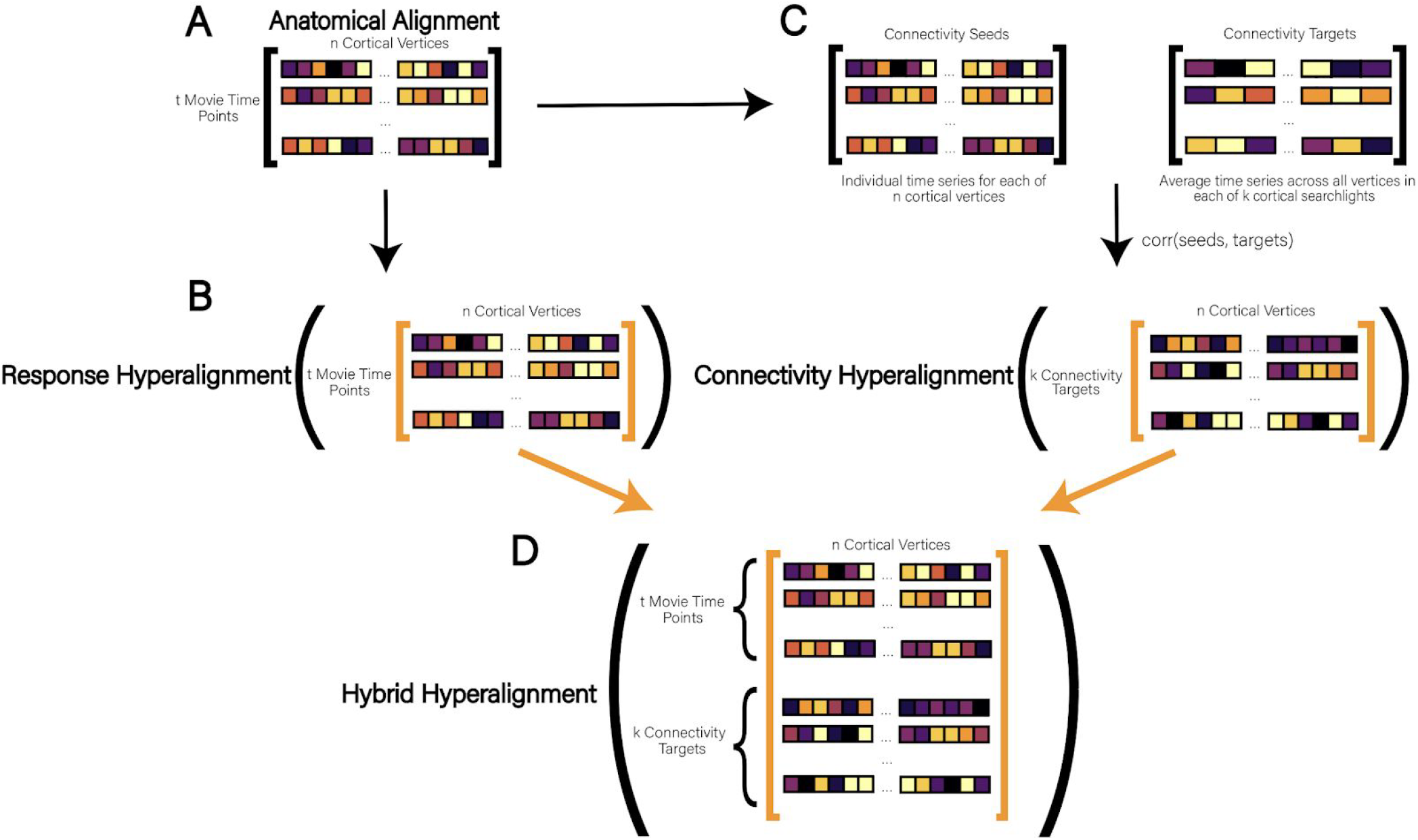
**(A)** In Anatomical Alignment, or “AA”, response profiles are aligned to a common anatomical template with *t* movie time points as rows and *n* cortical vertices as columns. **(B)** To perform Response Hyperalignment, AA data are passed directly to the searchlight hyperalignment algorithm to derive transformation matrices based solely on local activation-based activation profiles. **(C)** In Connectivity Hyperalignment, the time series of each cortical vertex is correlated with the average time series of vertices aggregated into larger targets across the brain (here, 1,076 searchlights). The resulting connectome with *k* connectivity targets as rows and *n* cortical vertices as columns is passed to searchlight hyperalignment to derive transformation matrices based solely on brain-wide connectivity profiles. **(D)** In our new method, Hybrid Hyperalignment, the data that would be used separately in Response Hyperalignment and Connectivity Hyperalignment are combined, resulting in (*t* movie time points + *k* connectivity targets) rows and *n* cortical vertices as columns. This data is then passed to the searchlight hyperalignment algorithm to derive transformations based on both local response and brain-wide connectivity profiles.

### 2.4.1 Response-Based Hyperalignment

To perform response-based hyperalignment we began with the AA data consisting of responses across cortical vertices (over time) in the downsampled fsaverage surface ("icoorder5", 3 mm resolution). We removed vertices within the medial wall for this analysis, which resulted in 9,372 and 9,370 vertices remaining in the left and right hemispheres respectively. The resulting data matrix for each participant consisted of a row for each TR (response patterns) and a column (18,742 total combined across left and right hemispheres) for each cortical surface vertex (Fig. 1B). Each time-series (column) of the matrix was z-scored to have zero mean and unit variance. This data served as input to the searchlight hyperalignment algorithm, which utilizes the Procrustes transformation to calculate a transformation matrix for each participant capable of mapping their AA data into a single high-dimensional information space shared across participants (Guntupalli et al., 2016).

The searchlight hyperalignment algorithm centers a searchlight on each cortical surface vertex and computes a common information space across participants for each searchlight. Because searchlights are highly overlapping, each vertex will be assigned transformation weights from multiple searchlight transformation matrices (Haxby et al., 2020). These transformation weights are aggregated and z-scored for each vertex to produce a single, whole-brain transformation matrix for each participant, which maps AA data into a single common space. The use of searchlights serves to constrain the Procrustes transformations of response profiles to a neuroanatomically meaningful radius. In other words, a vertex in the occipital lobe cannot be aligned to a vertex in the prefrontal cortex. Our analyses limited all Procrustes transformations to within a 20 mm searchlight radius (Guntupalli et al., 2016).

### 2.4.2 Connectivity-Based Hyperalignment

The implementation of connectivity-based hyperalignment is identical to that of RHA, except that CHA takes a connectivity data matrix as input, rather than a response data matrix. In a functional connectivity matrix, each row is a pattern of connectivity strengths across vertices (columns) for a “connectivity target” elsewhere in the brain. In this way, CHA distinguishes itself from RHA by functionally aligning brain data based on the co-activation of cortical vertices with the rest of the brain in contrast to using purely local response profiles.

To compute each participant’s connectomes (Fig. 1C), we began with the exact same data matrix as used as input to the RHA algorithm described above and then defined our connectivity seeds and targets. In this analysis, our connectivity seeds were of the same resolution as our data: each seed was an icoorder5 surface vertex. Our connectivity targets were defined on a sparser surface for two main reasons. By downsampling to a lower resolution, we reduced the number of data points and increased computational efficiency. More notably, defining dense connectivity targets (for example, vertex-to-vertex) on anatomically aligned data yields poor time-locked correspondence across participants (as shown in Figure 3). By aggregating these targets into searchlights, we ensure more reliable seed-target correspondence, which the Procrustes algorithm assumes. We define the vertices at the center of each connectivity target as each vertex on the icoorder3 surface (yielding 588 and 587 vertices in the left and right hemispheres respectively after masking the medial wall). We then centered a 13 mm searchlight on each of these vertices and computed an average time-series for each searchlight, which served as a connectivity target. We calculated the participant’s connectome as the correlation between the average time-series of each searchlight (connectivity targets) and the time-series of each icoorder5 vertex (connectivity seeds). Each column of a subject’s connectome was then z-scored to have zero-mean and unit variance, and the connectomes were passed to the searchlight hyperalignment algorithm in exactly the same process described above for response patterns in RHA. This produced a transformation matrix for each participant, which served to map each participant’s connectome (derived in AA space) into the newly derived connectivity-based common information space.

### 2.4.3 Hybrid Hyperalignment

The hybrid hyperalignment method combines both the neural response data inputted to RHA and the connectome inputted to CHA. The two data matrices do not necessarily have the same number of samples, as the samples of the response data represent the number of TRs collected and the samples of the connectome represent the number of connectivity targets we defined.Though each column in both of these matrices already had zero mean and unit variance, we wanted to ensure that the overall magnitudes of the variance of both RHA and CHA input data were the same, such that both types of information (response- and connectivity-based) would be equally weighted by the Procrustes transformation. We therefore applied a multiplier to every element of whichever input matrix contained fewer rows. To determine the multiplier, we calculated the Frobenius norm of both the response profile matrix and the connectome matrix for each participant. A ratio of the two Frobenius norms was then computed: the numerator of the ratio was the Frobenius norm of whichever input matrix contained more samples, and the denominator of the ratio was the Frobenius norm of whichever input matrix contained fewer samples.

Once this multiplier was applied, we vertically concatenated the connectome to the response data matrices (Fig. 1). The resulting matrix was of dimensions *t* time points plus 1,176 connectivity targets (rows/samples) by 18,742 vertices (columns/features). This matrix was then passed to the searchlight hyperalignment algorithm as described above with a 20 mm searchlight radius. Again, searchlight hyperalignment produced a transformation matrix for each participant capable of mapping their AA cortical data into a common information space, in this case based on both response and connectivity information.

It is important to note that all three hyperalignment methods made use of the same original neural data, but each method reoriented the dimensions of each individual’s anatomical space (cortical vertices) differently based on response pattern vectors only, connectivity pattern vectors only, or a weighted combination of response and connectivity pattern vectors.

## 2.5 Alignment Benchmarking

### 2.5.1 Intersubject Correlation of Response and Connectivity Profiles

To investigate the relative efficacy of the hyperalignment procedures in aligning shared information processing across brains, we computed the vertex-by-vertex intersubject correlation (Nastase et al., 2019) of both movie-viewing response profiles (time-series responses) (Fig. 2) and functional connectivity profiles (dense functional connectomes) (Guntupalli et al., 2018) (Fig. 3). First, the transformation matrices for each participant were calculated by RHA, CHA, and H2A separately using a leave-one-run-out data folding scheme described below. Next, participants’ held-out movie-viewing response profiles (test data) were mapped from anatomical space (fsaverage5) into each common space (derived from training data). Within anatomical space and each common space a dense, vertex-by-vertex functional connectome was computed by correlating each cortical vertex’s response time series with all 18,741 other vertices’ time series for every participant. The Pearson correlation was then calculated across participants for every vertex on both (1) the held-out response profile data and (2) the held-out dense functional connectomes in each of the 3 common information spaces. Differences in the distributions of ISCs across alignment algorithms were tested using a one-sided permutation test for AA vs. each hyperalignment method or a two-sided permutation test for comparing hyperalignment methods to each other (null distributions were created by shuffling alignment method labels 10,000 times in all tests). Mean ISCs across vertices, participants, and data folds were projected onto the fsaverage template for visualization.

**Figure 2:**
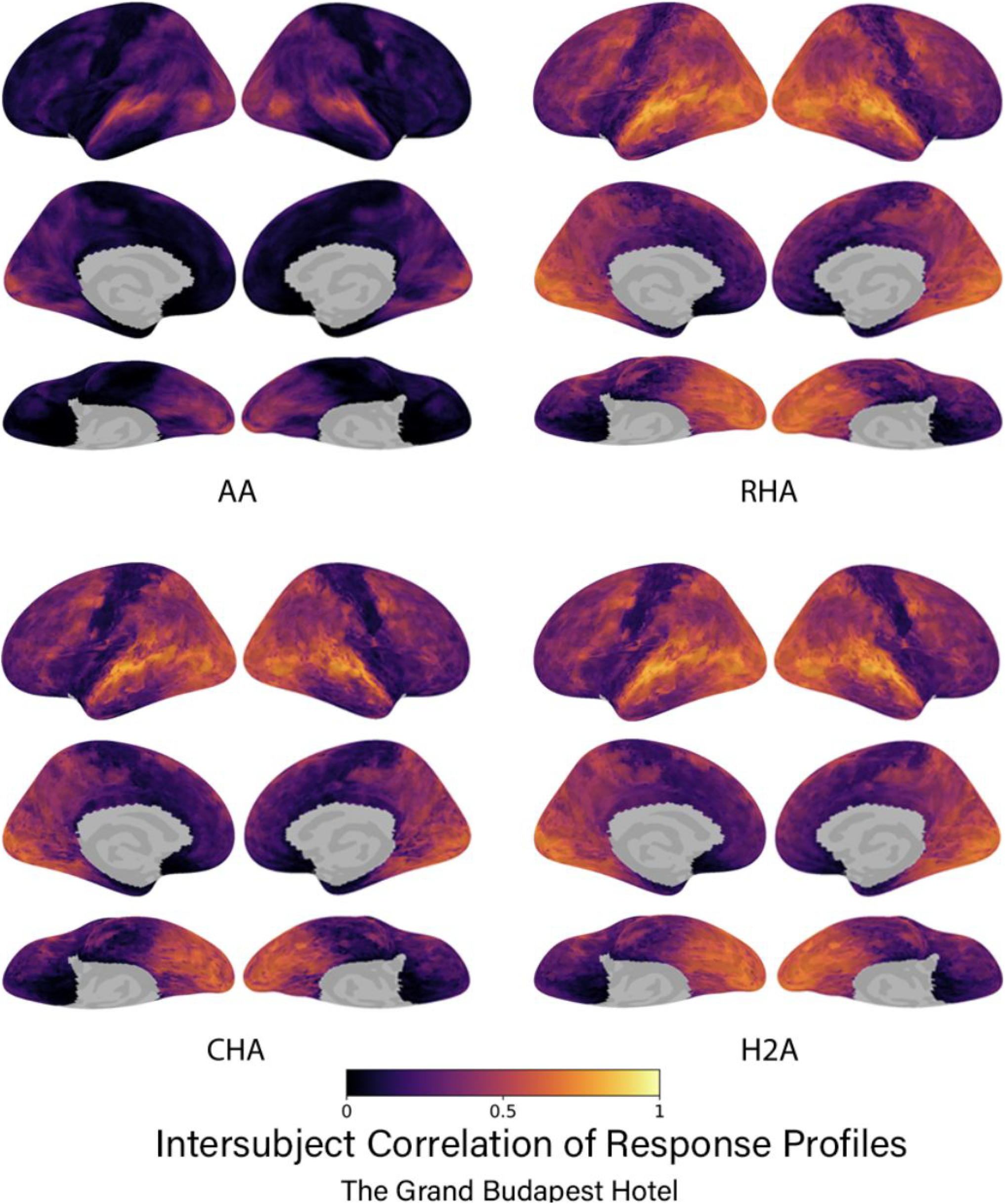
The intersubject correlation of response profiles using the Budapest data for each type of alignment algorithm. Correlations are presented for each vertex on the cortical surface averaged over data folds and participants. Subsequent figures show only left lateral hemisphere views of results. Brain image figures of results for all three datasets with lateral, medial, and ventral views are shown in Supplemental Figures S1 - S2.

**Figure 3:**
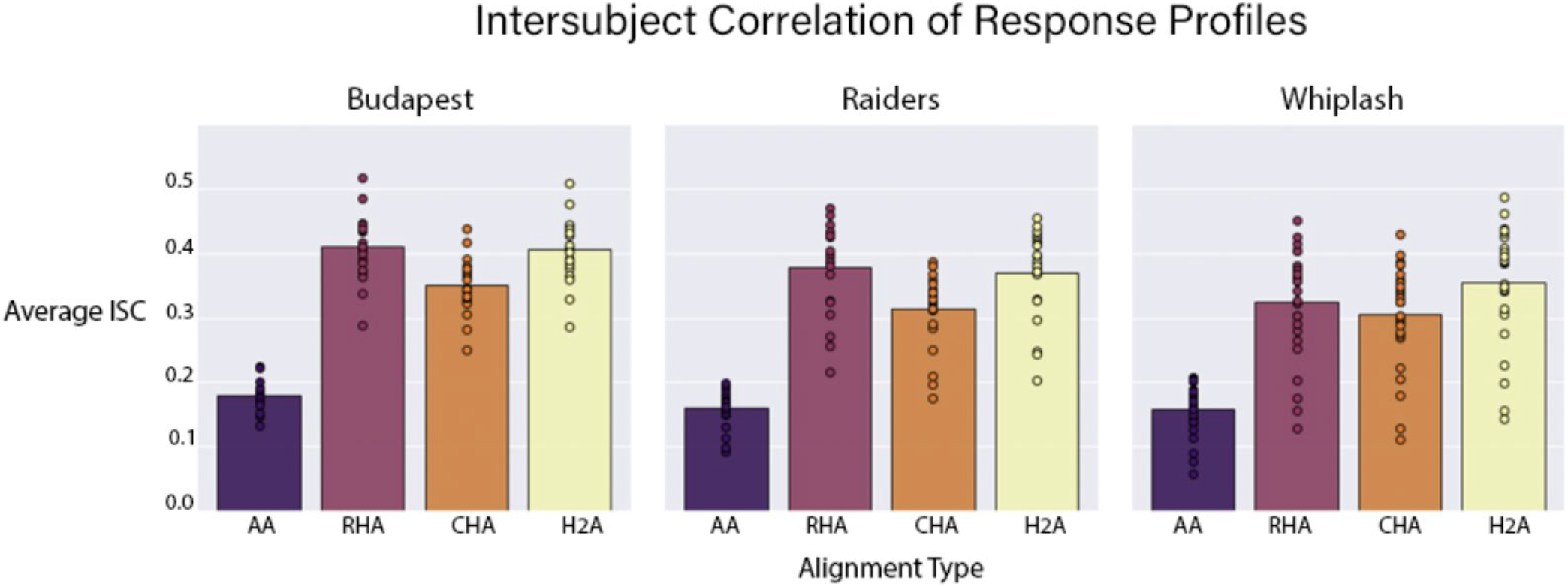
The average intersubject correlation of response profiles is shown for each alignment algorithm for each data set. Bars represent the average intersubject correlation over all vertices, data folds, and participants. Circles represent the average intersubject correlation for an individual participant over all vertices and data folds.

### 2.5.2 Movie Segment Classification

We computed the classification accuracies, searchlight-by-searchlight, of 5-second movie segments that were not used in the hyperalignment procedure. To do this, we compared each searchlight’s activity pattern (averaged across all vertices within a searchlight) in one participant with the average activity pattern over all other participants in the same searchlight for every 5-second movie segment (5 TRs). Ten-second buffer periods were added to both ends of every target segment such that no target segment was compared to a time segment within 10 seconds of itself.

The searchlights used for movie segment classification were centered on each cortical vertex and included all other vertices within a 13 mm radius of the center vertex. If a participant’s searchlight pattern of activation for a given segment was *most* similar to the group average response *for the corresponding segment* (relative to average group patterns for *all other* movie segments) it was considered correctly classified. We quantified “most similar” as the segment with the highest Pearson’s correlation coefficient. Differences in the distributions of accuracies for each subject across alignment algorithms were tested using a one-tailed permutation test for AA vs. each hyperalignment method or a two-tailed permutation test for comparing hyperalignment methods to each other. Null distributions were simulated by shuffling alignment method labels 10,000 times in all tests. Mean classification accuracies across searchlights, participants, and data folds were projected onto the fsaverage template for visualization.

### 2.5.3 Data Folding

We used a leave-one-run-out data folding scheme to validate hyperalignment training on an unseen portion of data. For each subject, hyperalignment parameters for each subject were trained on all but one run, and the held-out run was mapped into the trained space using the derived transformation matrix. Unseen data was mapped into the common model and alignment performance was benchmarked using our three chosen tests of intersubject alignment: response profile ISC, dense connectome ISC, and movie segment classification. ISC and classification analyses were therefore iteratively performed on every run of every movie after deriving a common space from all other runs from the same movie. Correlations and classification accuracies are reported as the average of these measures across data folds for each movie.

## 3 Results

### 3.1 Intersubject Correlation

#### 3.1.2 Response Profiles

All three hyperalignment algorithms in all three data sets yielded significant improvements in intersubject correlation of vertex time-series response profiles across participants relative to AA alone (p < 0.001 for all). Further, H2A aligned response profiles as well as RHA in all three data sets. In the Budapest data set, AA produced an average ISC of 0.179, while RHA, CHA, and H2A produced ISCs of 0.408, 0.349, and 0.406 respectively (Fig. 2A, B). RHA and H2A aligned response profiles significantly better than CHA (p < 0.001 for both), but were not significantly different from each other (p > 0.99). In the Raiders data set, AA produced an average ISC of 0.160, while RHA, CHA, and H2A yielded ISCs of 0.378, 0.314, and 0.370 respectively (Fig.2B). Again, RHA and H2A significantly outperformed CHA (p < 0.001), but were not significantly different from each other (p > 0.99). Finally, in the Whiplash data set, AA produced an average ISC of 0.158, while RHA, CHA, and H2A produced ISCs of 0.325, 0.306, and 0.354 respectively (Fig. 2B). In this dataset RHA and H2A performed significantly better than CHA (p < 0.001 for both), and H2A performed significantly better than RHA (p < 0.001). Of note, the Whiplash data set was only about half the duration of the other two data sets, which may partially account for why the ISCs across alignment methodologies are lower for these participants.

#### 3.1.2 Dense Connectivity Profiles

All three hyperalignment procedures significantly improved the intersubject alignment of dense connectivity profiles relative to AA alone across data sets (p < 0.001 for all), with H2A consistently producing the highest ISCs of any method. In the Budapest data set, AA produced an average ISC of 0.437, while RHA, CHA, and H2A produced ISCs of 0.793, 0.807, and 0.857 respectively (Fig 3A, B). The ISCs of CHA and RHA were not significantly different (p = 0.850), but the ISC of H2A was significantly higher than both CHA and RHA (p < 0.001 for both). When aligning on the Raiders data, AA produced an average ISC of 0.417, and RHA, CHA, and H2A yielded ISCs of 0.762, 0.789, and 0.847 respectively (Fig. 3B). Again, the ISCs of CHA and RHA were not significantly different (p = 0.987), but the ISC of H2A was significantly higher than both CHA and RHA (p < 0.001 for both). Finally, in the smaller Whiplash data set, AA had an average ISC of 0.285, and RHA, CHA, and H2A resulted in ISCs of 0.566, 0.618, and 0.675 respectively (Fig. 3B). In this data, the ISCs of both CHA and H2A were significantly greater than RHA (p < 0.001 for both). Further, the ISCs of H2A were significantly greater than those of CHA. The shorter duration of the Whiplash movie-viewing session may partially account for the lower ISCs across alignment algorithms.

**Figure 4:**
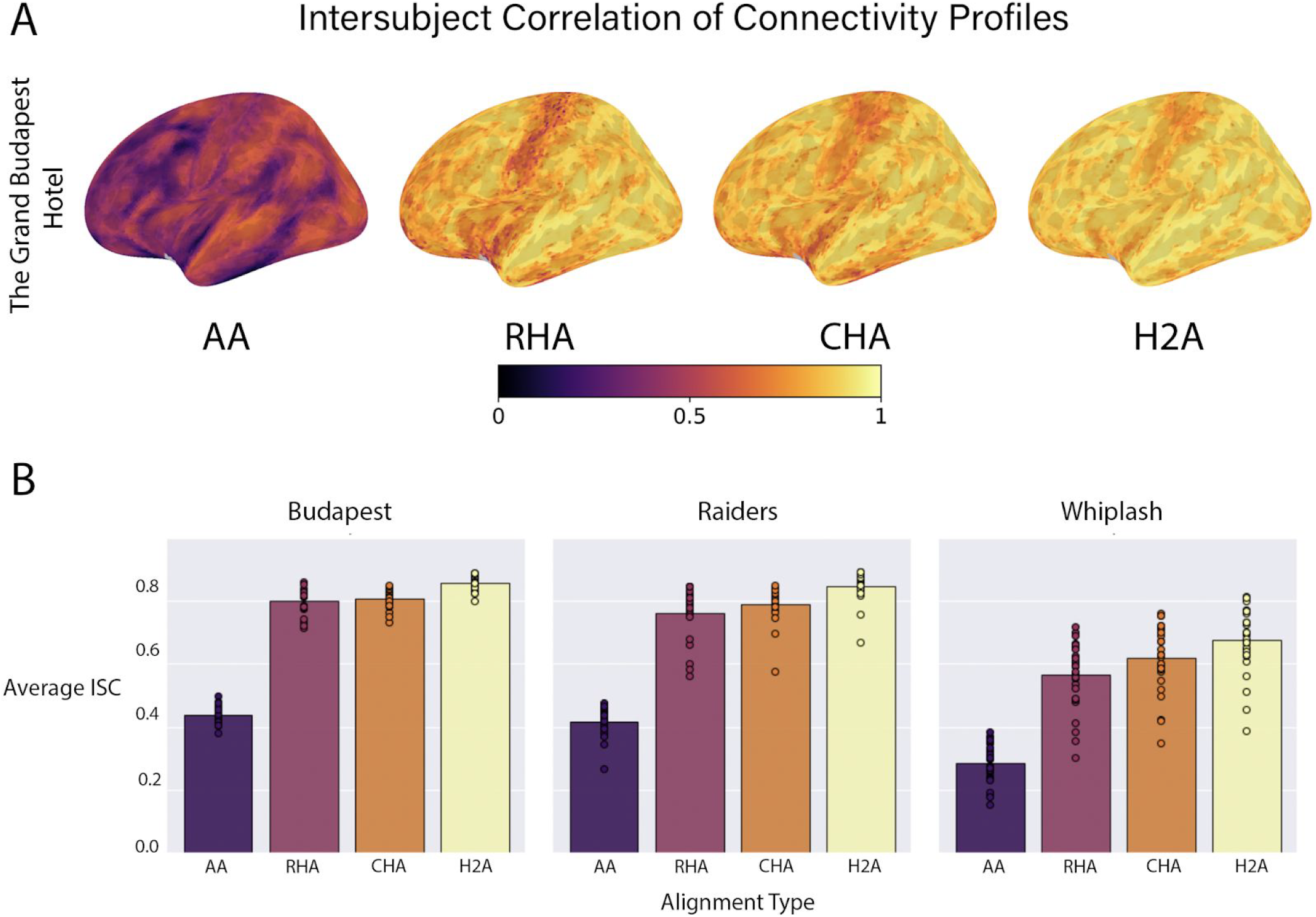
The average intersubject correlation of connectivity profiles. (A) Correlations are presented for each vertex on the left lateral cortical surface averaged over data folds and participants. Brain image figures of results with lateral, medial, and ventral views of both hemispheres are shown in Supplemental Figures S3 - S5. (B) Correlations are shown for each alignment algorithm for each data set. Bars represent the average intersubject correlation over all vertices, data folds, and participants. Circles represent the average intersubject correlation for an individual participant over all vertices and data folds.

### 3.2 Movie Segment Classification

Hyperalignment, regardless of the specific algorithm, showed significant improvements relative to AA in classifying 5-second movie segments (p < 0.001 for all). In nearly every common space across data sets, the individual with the *lowest* hyperaligned classification accuracy had better accuracy than the individual with the *highest* AA accuracy (Fig. 4B). We present results here as the average classification accuracy across searchlights, participants, and data folds. In the Budapest data set, AA produced an average accuracy of 0.020, while RHA, CHA, and H2A had accuracies of 0.165, 0.115, and 0.145 respectively (Fig. 4A). In this data set, RHA and H2A both classified time segments better than CHA (p < 0.001 for both), and RHA significantly outperformed H2A (p < 0.001). In the Raiders data set, AA produced an average classification accuracy of 0.012, and RHA, CHA, and H2A yielded accuracies of 0.118, 0.076, and 0.100 respectively. Again, RHA and H2A were both significantly better than CHA at classifying time segments (p < 0.001 for both), and RHA significantly outperformed H2A (p < 0.001). Finally, in the Whiplash data set, AA had an average accuracy of 0.019, while RHA, CHA, and H2A produced accuracies of 0.136, 0.104, and 0.119. In this data set RHA and H2A significantly outperformed CHA (p < 0.001 for both), and again, RHA significantly outperformed H2A (p < 0.001).

**Figure 5:**
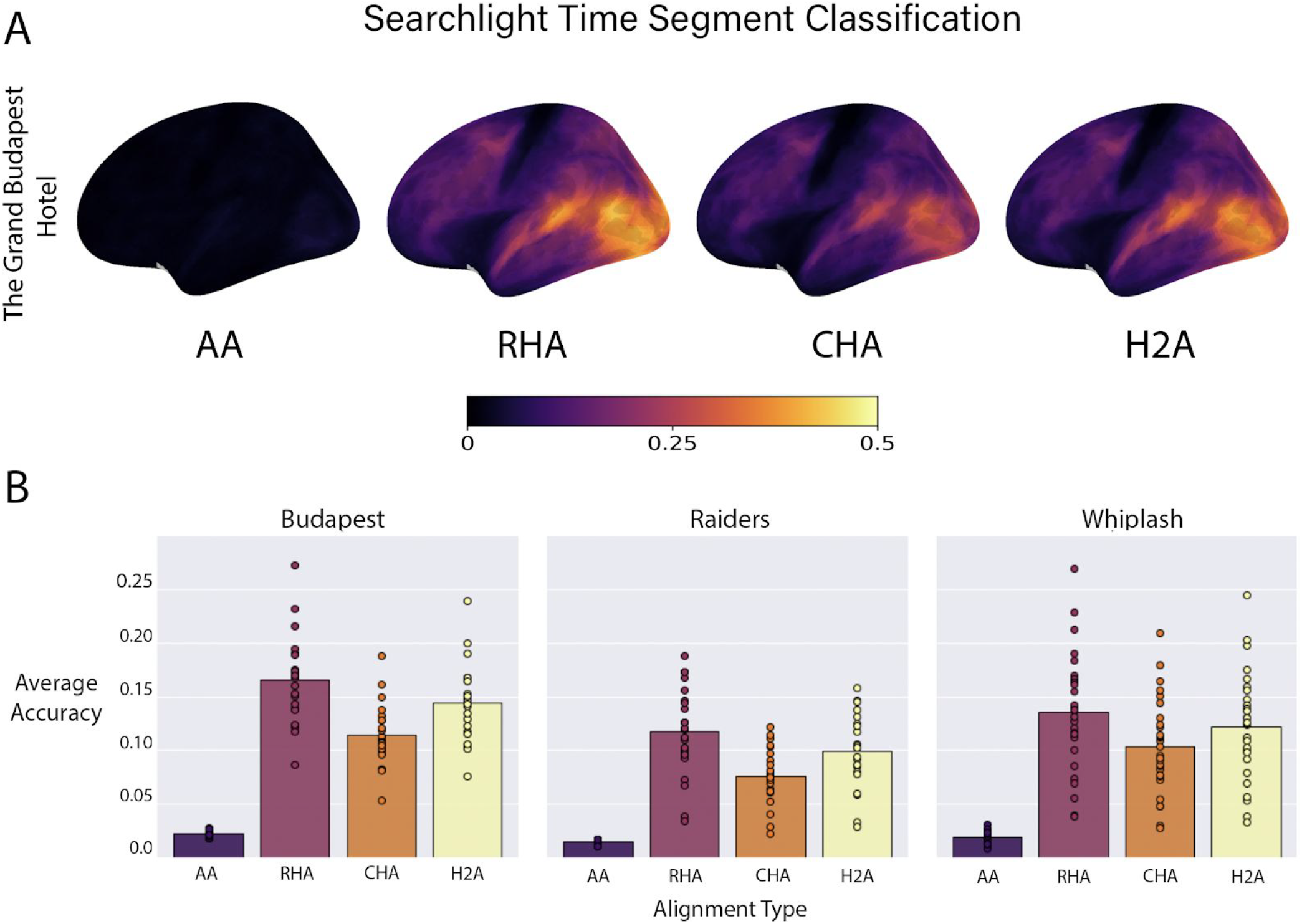
Average time segment classification accuracies. (A) Accuracies are presented for all searchlights on the left lateral cortical surface averaged over data folds and participants. Brain image figures of results with lateral, medial, and ventral views of both hemispheres are shown in Supplemental Figures S6 - S8. (B) Correlations are shown for each alignment algorithm for each data set. Bars represent the average classification accuracies over all searchlights, data folds, and participants. Circles represent the average classification accuracy for an individual participant over all vertices and data folds.

## 4 Discussion

A major objective of the hyperalignment algorithm is to map the shared information originally found in idiosyncratic cortical topographies into a common space in which this information is better aligned across participants. Previously, RHA was shown to align response-based data better than CHA, whereas CHA was shown to better align connectivity-based data than RHA. In this study we used three separate data sets to show that a hybrid hyperalignment algorithm, H2A, which uses both response and connectivity data from the same fMRI dataset to derive transformation matrices, is capable of aligning both types of data in a single common information space. Adding response information in the derivation of the common information space clearly improves the alignment of connectivity information. Adding connectivity information marginally improved alignment of response information on one measure - ISC of response profiles - but slightly degraded performance on another - bsMVPC of movie time segments.

H2A showed nearly identical improvements in the ISC of response profiles to those of RHA across data sets. In the Budapest and Raiders data sets H2A produced ISCs of response profiles that were as large as those produced by RHA (Fig. 2). In the Whiplash data set H2A outperformed RHA in aligning response profiles across participants. These results show that the new hybrid hyperalignment algorithm derives a common space that is capable of aligning shared response profiles as well as or better than RHA.

In addition to aligning response-based information, H2A models aligned connectivity-based information in the same common space. For all three data sets, H2A yielded improvements in the ISC of connectivity profiles that were greater than that of CHA (Fig. 3). H2A shows that the combination of response- and connectivity-based information for deriving the common model significantly improves the alignment of these connectomes. Response patterns provide additional information that helps to fine-tune the parameters in transformation matrices for alignment of connectivity patterns.

Finally, in the strictest test of the alignment of cortex-wide response patterns, we classified 5-second movie time segments by comparing each individual’s response pattern to the average group response pattern (See Movie Segment Classification above). In this analysis we found that H2A performed nearly as well as RHA. Permutation tests showed RHA performing significantly better than H2A, with only small, nearly identical differences between the two algorithms’ classification accuracies across the three datasets, ranging from 0.017 to 0.020.

Our findings indicate that functional alignment based upon either response or functional connectivity information alone provides an imperfect estimate of an optimal common space that maximizes the shared information we can account for between brains. Combining response and connectivity information to derive individual transformation matrices differentially impacted the intersubject alignment of response profiles and connectivity profiles. The addition of response information greatly improved alignment of connectivity profiles (Fig. 3), which showed significantly greater ISCs of dense connectomes for data aligned using H2A relative to CHA. At the same time, the addition of connectivity information did not consistently improve the alignment of response profiles. Further, forcing the algorithm to align connectivity profiles as well as response profiles in H2A slightly but significantly degraded between-subject movie segment classifications, our most stringent test of response alignment. This asymmetry suggests that the information provided by connectivity pattern vectors may not be as powerful as the information provided by response pattern vectors for fine-tuning the parameters in the transformation matrices.

We suspect that our time segment classification results reveal that the connectome we used in H2A may have lost information that would help to fine-tune the parameters in hyperalignment transformation matrices. This could be explained by our coarse definition of connectivity targets as mean time-series for large cortical fields rather than by an intrinsic weakness in using the information embedded within connectivity profiles to derive transformation matrices. At the same time, increasing the granularity of connectivity targets is not trivial because finer targets have decreased correspondence across participants. Increasing fine-grained information in the connectome may require noise reduction algorithms like principal component analysis to increase correspondence across participants (Guntupalli et al., 2018) or partial least squares analysis (Wold et al., 2001; Krishnan et al., 2011) to derive more meaningful time-series from multivariate patterns in each searchlight. It is possible that alternative ways to define the connectivity targets for hyperalignment could increase the information content in connectivity matrices and, thereby, increase the power of CHA and H2A. A more robust, functionally-meaningful connectome could mitigate the asymmetry in the relative utility of connectivity information versus response information.

Another other consideration is that we applied a multiplier to either the response or connectivity input data for H2A such that the Frobenius norms of both data matrices were equal. It is possible that unequal weighting of the two types of data may in fact be optimal for deriving H2A transformation matrices. For example, it may be preferable to weight RHA more heavily in visual areas and CHA more heavily in prefrontal areas. We plan to explore this idea further in future studies.

Despite H2A’s evident improvement in aligning functional connectomes compared with CHA, there are some intrinsic limitations that apply to H2A but not CHA. H2A and RHA both require that participants share the same time-locked stimulus with the same number of time points, so they cannot be applied to resting-state data or data sets that implement different stimuli.Because CHA aligns functional connectivity profiles rather than time series data, it alone can be used with datasets that don’t have time-locked stimuli (Nastase et al., 2020).

In comparison to other methods of functional alignment, our novel H2A method aligns both response and connectivity information using a single algorithm. Many researchers are interested in discerning both specific vertex-wise patterns of activation and patterns of functional network connectivity that correspond to different states of consciousness. Previously, fully leveraging hyperalignment to conduct both of these types of analyses would require implementing RHA for investigating response patterns and CHA separately for investigating functional connectivity.With the new H2A algorithm, researchers can run hyperalignment once and use the single hybrid common space to address both response and connectivity-oriented questions.

## 5 Conclusions

Our results suggest that there exists a single common information space capable of modeling shared response and connectivity information between brains. If optimization of shared response and connectivity information resulted in two separate common spaces, the derivation of a single common space using both types of information should vitiate its alignment capabilities. Instead, we found that a hybrid common space aligns response data as well as RHA and connectivity data better than CHA. This suggests that the two methods individually produce imperfect estimates of a single optimal information space. The H2A algorithm capitalizes on the strengths of different types of information to provide a more robust estimate of this optimal information space. This makes the H2A algorithm a preferable method for aligning stimulus response data when one wants to evaluate both connectivity and response data.However, H2A does require data collected while participants are shown a time-locked stimulus such as a movie. In cases where this type of data is unavailable, CHA can still be used to align shared information. Our new single-procedure algorithm is a powerful tool for elucidating the underlying space that encodes various forms of information represented in the brain.

## Acknowledgements

We thank Yaroslav O. Halchenko for helpful discussions and software advice. We also thank John Hudson from Dartmouth Research Computing for support with feisty environments and conflicting multiprocessing packages.

## Data/Code Availability

The *Grand Budapest Hotel* data are publicly available on *Scientific Data* (Visconti di Oleggio Castello, Chauhan, et al., 2020). The *Raiders of the Lost Ark* and *Whiplash* data will be made available at time of publication.

Response and connectivity based hyperalignment implementations are available publicly as part of the PyMVPA software package (Hanke et al., 2009). The hybrid hyperalignment implementation will be made available via PyMVPA, and all analysis code will be released as well.

## Contributions

**Erica L. Busch**: Conceptualization, methodology, software, validation, formal analysis, investigation, data curation, writing - reviewing and editing, visualization. **Lukas Slipski**: Conceptualization, methodology, validation, formal analysis, investigation, data curation, writing - original draft, visualization. **Ma Feilong**: Conceptualization, methodology, validation, supervision, writing - reviewing and editing. **J. Swaroop Guntapalli**: Software, data curation, writing - reviewing and editing. **Matteo Visconti di Oleggio Castello**: Data curation, writing - reviewing and editing. **Jeremy F. Huckins**: Data curation, writing - reviewing and editing.**Samuel A. Nastase**: Data curation, writing - reviewing and editing. **M. Ida Gobbini**: Data curation, funding acquisition. **Tor D. Wager**: Methodology, validation, writing - reviewing and editing. **James V. Haxby**: Conceptualization, methodology, validation, investigation, resources, writing - reviewing and editing, supervision, funding acquisition.

