## Supplemental Materials for "Hybrid Hyperalignment: A single high-dimensional model of shared information embedded in cortical patterns of response and functional connectivity"

### Supplemental figures

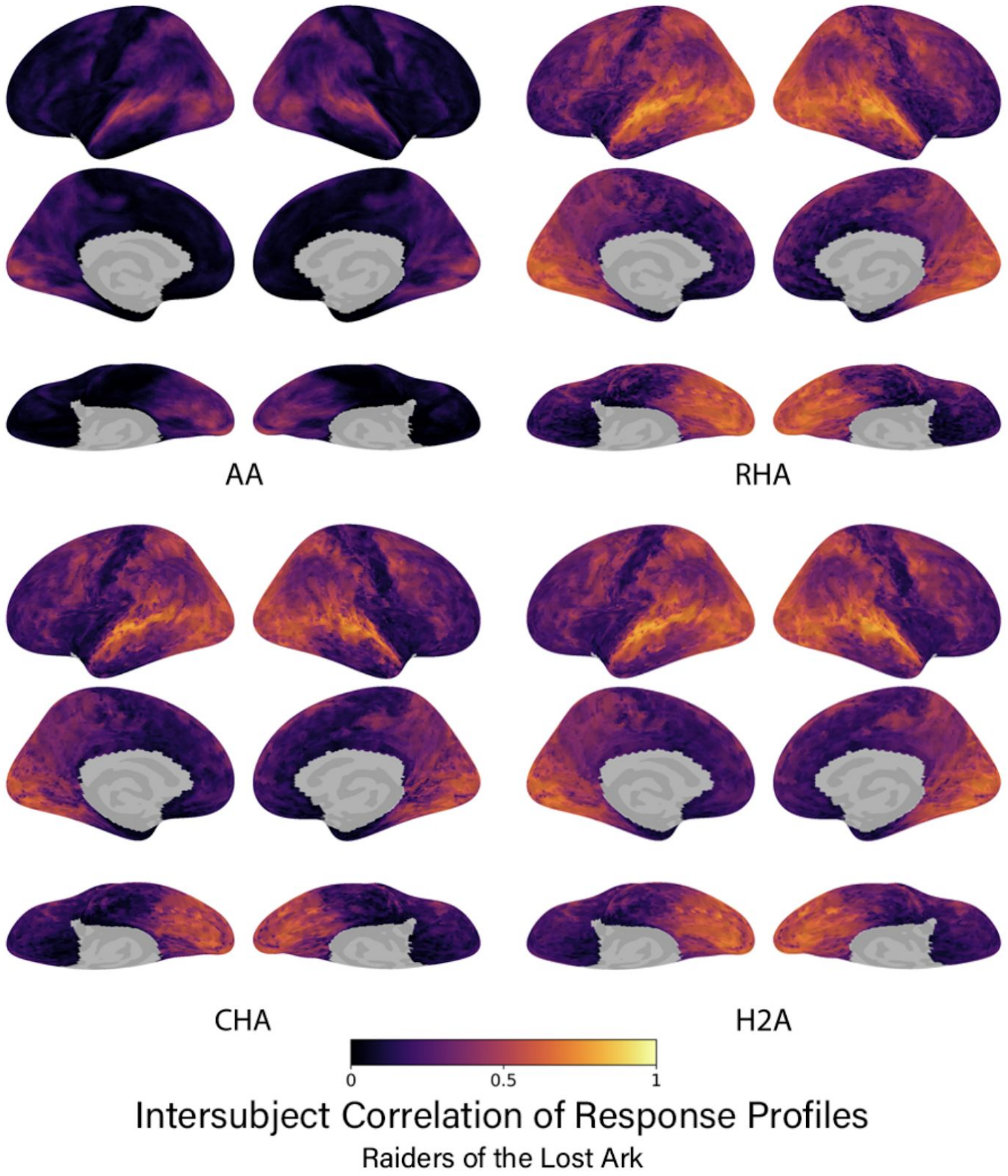


**S1.** The intersubject correlation of response profiles using the Raiders data for each type of alignment algorithm. Correlations are presented for each vertex on the cortical surface averaged over data folds and participants. Lateral, medial, and ventral views shown.

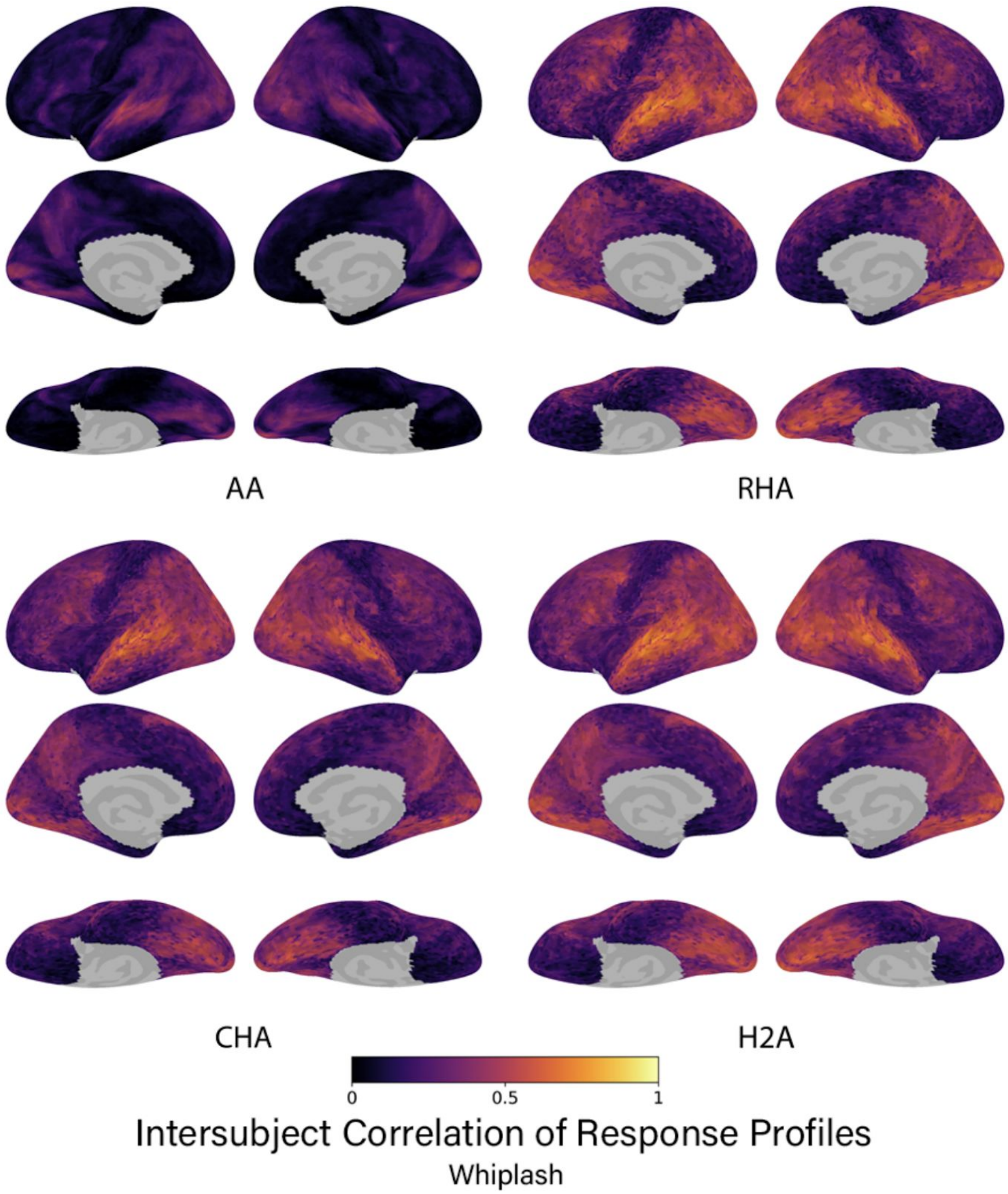


**S2.** The intersubject correlation of response profiles using the Whiplash data for each type of alignment algorithm. Correlations are presented for each vertex on the cortical surface averaged over data folds and participants. Lateral, medial, and ventral views shown.

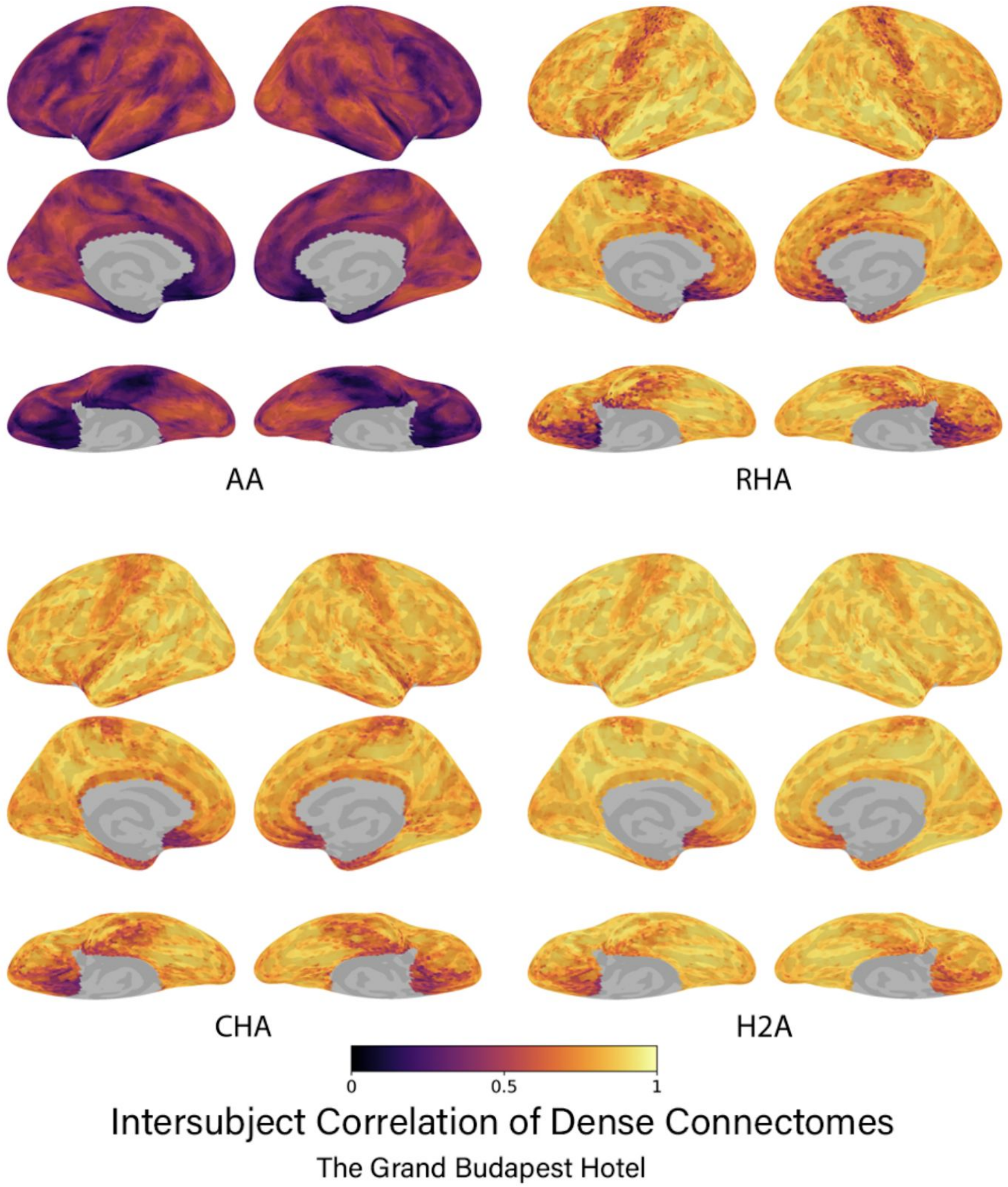


**S3.** The intersubject correlation of connectivity profiles using the Budapest data for each type of alignment algorithm. Correlations are presented for each vertex on the cortical surface averaged over data folds and participants. Lateral, medial, and ventral views shown.

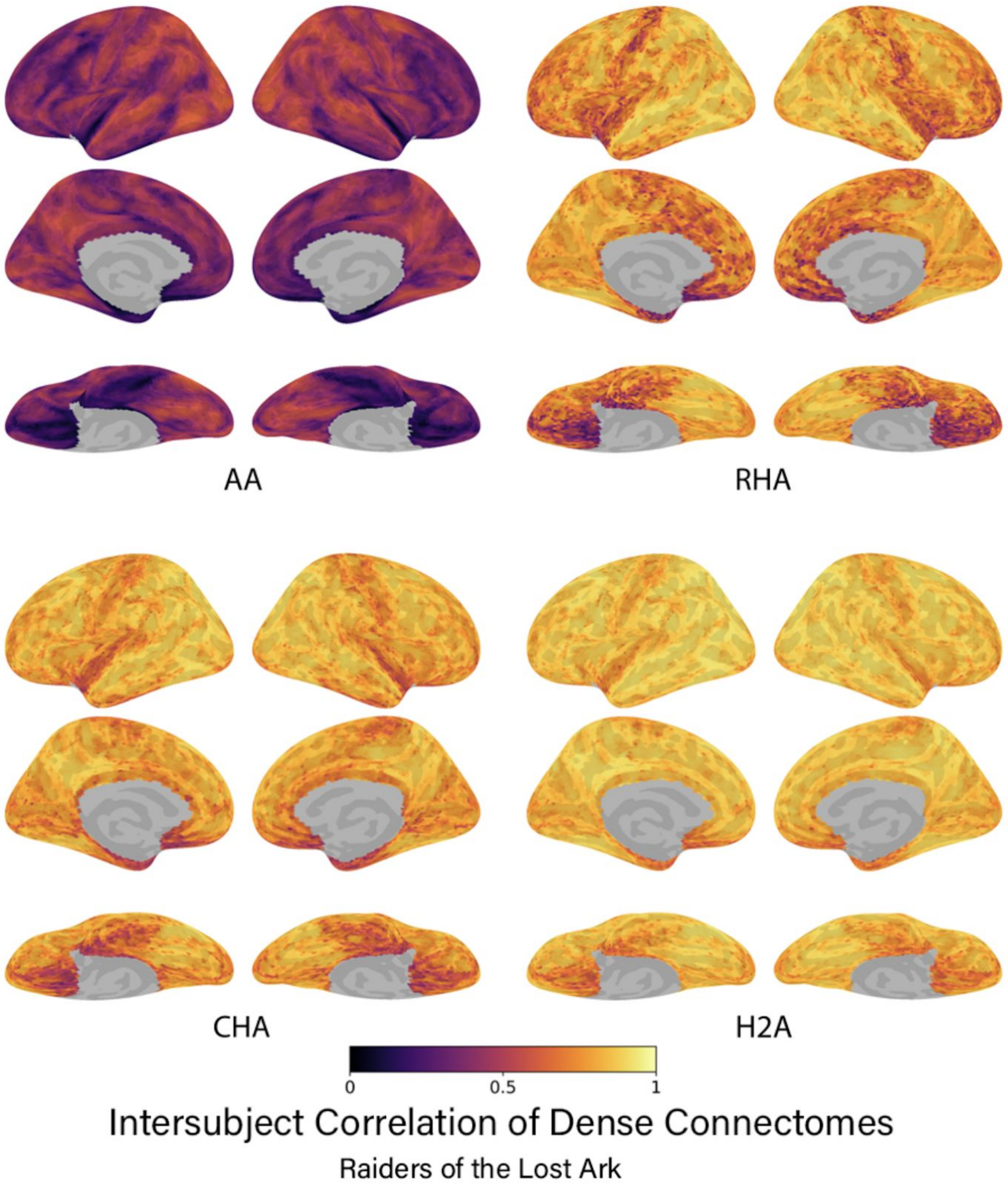


**S4.** The intersubject correlation of connectivity profiles using the Raiders data for each type of alignment algorithm. Correlations are presented for each vertex on the cortical surface averaged over data folds and participants. Lateral, medial, and ventral views shown.

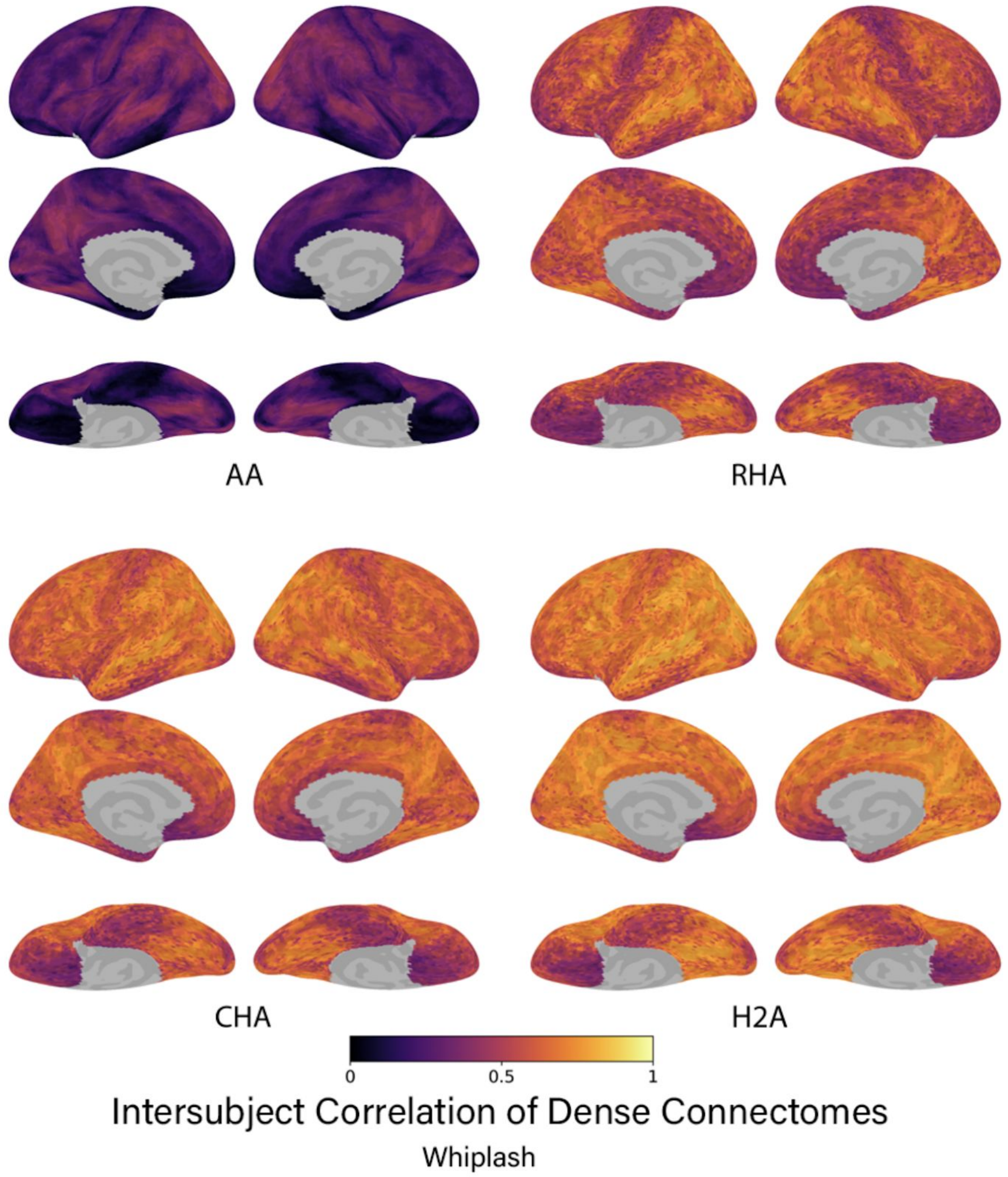


**S5.** The intersubject correlation of connectivity profiles using the Whiplash data for each type of alignment algorithm. Correlations are presented for each vertex on the cortical surface averaged over data folds and participants. Lateral, medial, and ventral views shown.

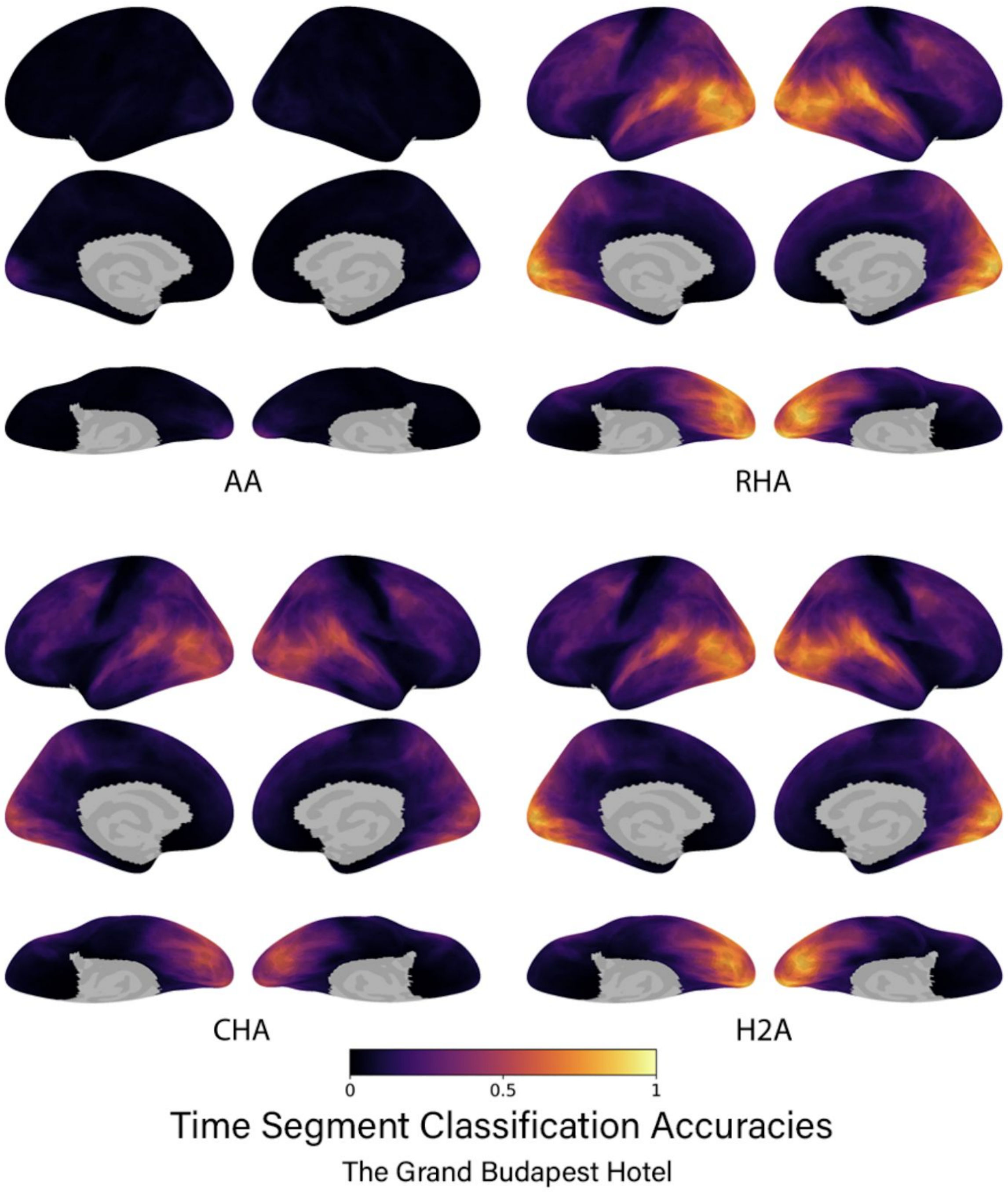


**S6.** The time segment classification accuracies using the Budapest data for each type of alignment algorithm. Accuracies are presented for all searchlights on the cortical surface averaged over data folds and participants. Lateral, medial, and ventral views shown.

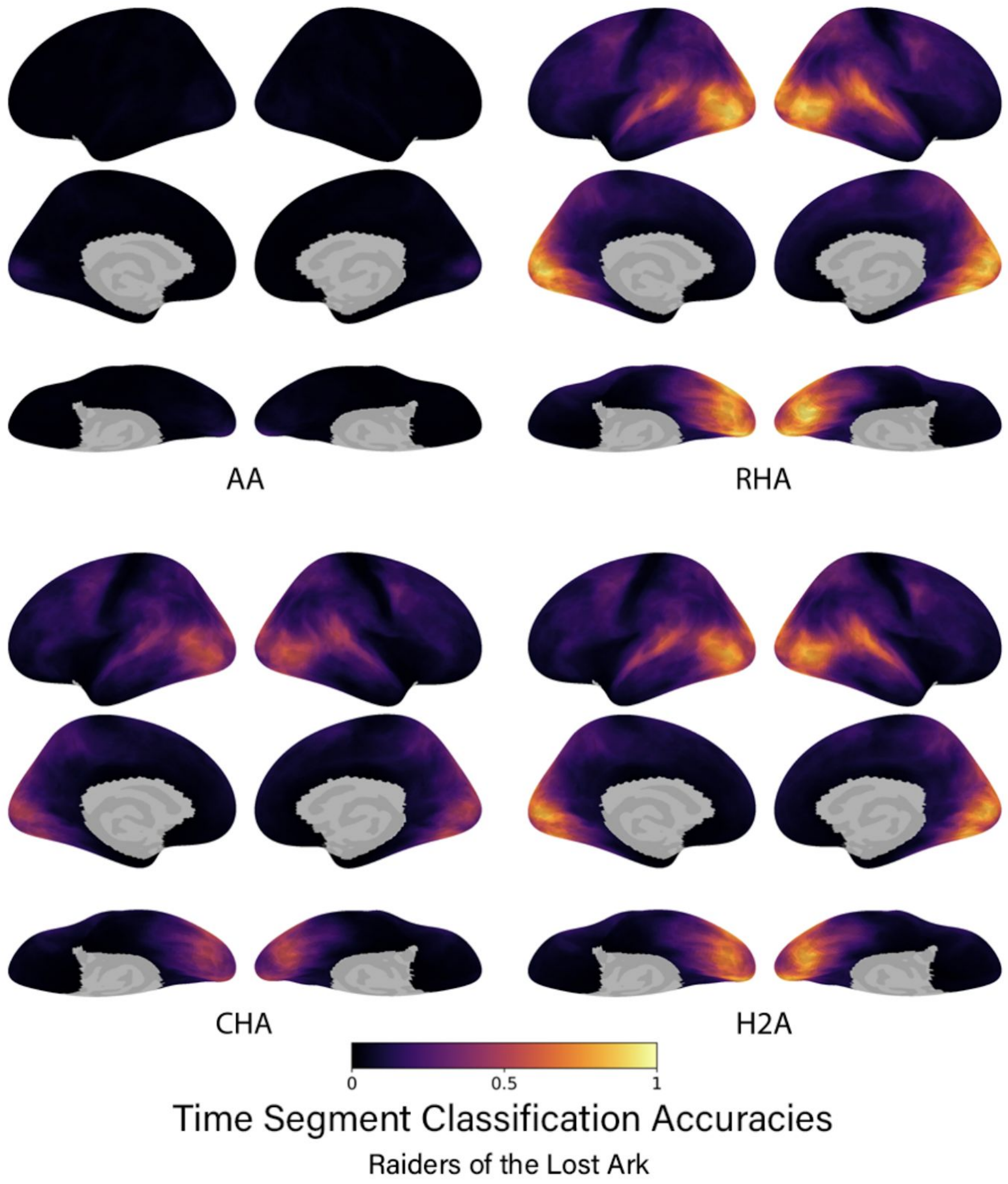


**S7.** The time segment classification accuracies using the Raiders data for each type of alignment algorithm. Accuracies are presented for all searchlights on the cortical surface averaged over data folds and participants. Lateral, medial, and ventral views shown.

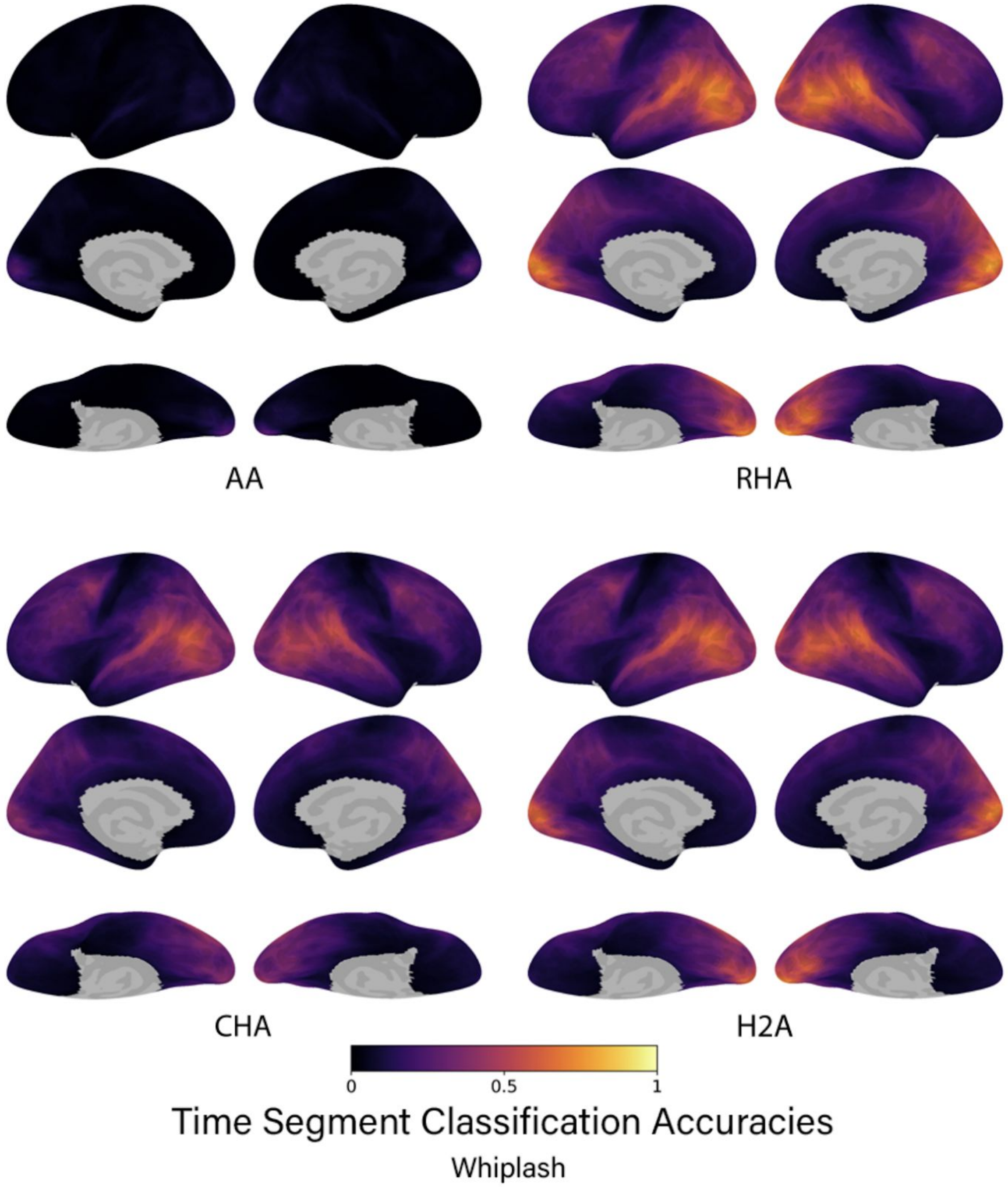


**S8.** The time segment classification accuracies using the Whiplash data for each type of alignment algorithm. Accuracies are presented for all searchlights on the cortical surface averaged over data folds and participants. Lateral, medial, and ventral views shown.

### Supplemental methods

#### Preprocessing Parameters

Whiplash results included in this manuscript come from preprocessing performed using *fMRIPrep* 20.0.3 (Esteban, Markiewicz, et al. (2018); Esteban, Blair, et al. (2018); RRID:SCR_016216), which is based on *Nipype* 1.4.2 (Gorgolewski et al. (2011); Gorgolewski et al. (2018); RRID:SCR_002502). Budapest and Raiders results used *fMRIPrep 20.1.1.*

#### Anatomical data preprocessing

A total of 3 T1-weighted (T1w) images were found within the input BIDS dataset. All of them were corrected for intensity non-uniformity (INU) with N4BiasFieldCorrection (Tustison et al. 2010), distributed with ANTs 2.2.0 (Avants et al. 2008, RRID:SCR_004757). The T1w-reference was then skull-stripped with a *Nipype* implementation of the antsBrainExtraction.sh workflow (from ANTs), using OASIS30ANTs as target template. Brain tissue segmentation of cerebrospinal fluid (CSF), white-matter (WM) and gray-matter (GM) was performed on the brain-extracted T1w using fast (FSL 5.0.9, RRID:SCR_002823, Zhang, Brady, and Smith 2001). A T1w-reference map was computed after registration of 3 T1w images (after INU-correction) using mri_robust_template (FreeSurfer 6.0.1, Reuter, Rosas, and Fischl 2010). Brain surfaces were reconstructed using recon-all (FreeSurfer 6.0.1, RRID:SCR_001847, Dale, Fischl, and Sereno 1999), and the brain mask estimated previously was refined with a custom variation of the method to reconcile ANTs-derived and FreeSurfer-derived segmentations of the cortical gray-matter of Mindboggle (RRID:SCR_002438, Klein et al. 2017). Volume-based spatial normalization to two standard spaces (MNI152NLin2009cAsym, MNI152NLin6Asym) was performed through nonlinear registration with antsRegistration (ANTs 2.2.0), using brain-extracted versions of both T1w reference and the T1w template. The following templates were selected for spatial normalization: *ICBM 152 Nonlinear Asymmetrical template version 2009c* [Fonov et al. (2009), RRID:SCR_008796; TemplateFlow ID: MNI152NLin2009cAsym], *FSL's MNI ICBM 152 non-linear 6th Generation Asymmetric Average Brain Stereotaxic Registration Model* [Evans et al. (2012), RRID:SCR_002823; TemplateFlow ID: MNI152NLin6Asym],

#### Functional data preprocessing

For each of the 15 BOLD runs found per subject (across all tasks and sessions), the following preprocessing was performed. First, a reference volume and its skull-stripped version were generated using a custom methodology of *fMRIPrep*. A deformation field to correct for susceptibility distortions was estimated based on *fMRIPrep*'s *fieldmap-less* approach. The deformation field is that resulting from co-registering the BOLD reference to the same-subject T1w-reference with its intensity inverted (Wang et al. 2017; Huntenburg 2014). Registration is performed with antsRegistration (ANTs 2.2.0), and the process regularized by constraining deformation to be nonzero only along the phase-encoding direction, and modulated with an average fieldmap template (Treiber et al. 2016). Based on the estimated susceptibility distortion, a corrected EPI (echo-planar imaging) reference was calculated for a more accurate co-registration with the anatomical reference. The BOLD reference was then co-registered to the T1w reference using bbregister (FreeSurfer) which implements boundary-based registration (Greve and Fischl 2009). Co-registration was configured with six degrees of freedom. Head-motion parameters with respect to the BOLD reference (transformation matrices, and six corresponding rotation and translation parameters) are estimated before any spatiotemporal filtering using mcflirt (FSL 5.0.9, Jenkinson et al. 2002). The BOLD time-series were resampled onto the following surfaces (FreeSurfer reconstruction nomenclature): *fsaverage*. The BOLD time-series (including slice-timing correction when applied) were resampled onto their original, native space by applying a single, composite transform to correct for head-motion and susceptibility distortions. These resampled BOLD time-series will be referred to as *preprocessed BOLD in original space*, or just *preprocessed BOLD*. The BOLD time-series were resampled into standard space, generating a *preprocessed BOLD run in MNI152NLin2009cAsym space*. First, a reference volume and its skull-stripped version were generated using a custom methodology of *fMRIPrep*. *Grayordinates* files (Glasser et al. 2013) containing 91k samples were also generated using the highest-resolution fsaverage as intermediate standardized surface space. Several confounding time-series were calculated based on the *preprocessed BOLD*: framewise displacement (FD), DVARS and three region-wise global signals. FD and DVARS are calculated for each functional run, both using their implementations in *Nipype* (following the definitions by Power et al. 2014). The three global signals are extracted within the CSF, the WM, and the whole-brain masks. Additionally, a set of physiological regressors were extracted to allow for component-based noise correction (*CompCor*, Behzadi et al. 2007). Principal components are estimated after high-pass filtering the *preprocessed BOLD* time-series (using a discrete cosine filter with 128s cut-off) for the two *CompCor* variants: temporal (tCompCor) and anatomical (aCompCor). tCompCor components are then calculated from the top 5% variable voxels within a mask covering the subcortical regions. This subcortical mask is obtained by heavily eroding the brain mask, which ensures it does not include cortical GM regions. For aCompCor, components are calculated within the intersection of the aforementioned mask and the union of CSF and WM masks calculated in T1w space, after their projection to the native space of each functional run (using the inverse BOLD-to-T1w transformation). Components are also calculated separately within the WM and CSF masks. For each CompCor decomposition, the *k* components with the largest singular values are retained, such that the retained components' time series are sufficient to explain 50 percent of variance across the nuisance mask (CSF, WM, combined, or temporal). The remaining components are dropped from consideration. The head-motion estimates calculated in the correction step were also placed within the corresponding confounds file. The confound time series derived from head motion estimates and global signals were expanded with the inclusion of temporal derivatives and quadratic terms for each (Satterthwaite et al. 2013). Frames that exceeded a threshold of 0.25 mm FD or 1.05 standardised DVARS were annotated as motion outliers. All resamplings can be performed with *a single interpolation step* by composing all the pertinent transformations (i.e. head-motion transform matrices, susceptibility distortion correction when available, and co-registrations to anatomical and output spaces). Gridded (volumetric) resamplings were performed using antsApplyTransforms (ANTs), configured with Lanczos interpolation to minimize the smoothing effects of other kernels (Lanczos 1964). Non-gridded (surface) resamplings were performed using mri_vol2surf (FreeSurfer).

Many internal operations of *fMRIPrep* use *Nilearn* 0.6.2 (Abraham et al. 2014, RRID:SCR_001362), mostly within the functional processing workflow. For more details of the pipeline, see the section corresponding to workflows in *fMRIPrep*'s documentation.

**Copyright Waiver**

The above boilerplate text was automatically generated by fMRIPrep with the express intention that users should copy and paste this text into their manuscripts *unchanged*. It is released under the CC0 license.

**References**

Abraham, Alexandre, Fabian Pedregosa, Michael Eickenberg, Philippe Gervais, Andreas Mueller, Jean Kossaifi, Alexandre Gramfort, Bertrand Thirion, and Gael Varoquaux. 2014.

“Machine Learning for Neuroimaging with Scikit-Learn.” *Frontiers in Neuroinformatics* 8. https://doi.org/10.3389/fninf.2014.00014.

Avants, B.B., C.L. Epstein, M. Grossman, and J.C. Gee. 2008. “Symmetric Diffeomorphic Image Registration with Cross-Correlation: Evaluating Automated Labeling of Elderly and Neurodegenerative Brain.” *Medical Image Analysis* 12 (1): 26–41. https://doi.org/10.1016/j.media.2007.06.004.

Behzadi, Yashar, Khaled Restom, Joy Liau, and Thomas T. Liu. 2007. “A Component Based Noise Correction Method (CompCor) for BOLD and Perfusion Based fMRI.” *NeuroImage* 37 (1): 90–101. https://doi.org/10.1016/j.neuroimage.2007.04.042.

Dale, Anders M., Bruce Fischl, and Martin I. Sereno. 1999. “Cortical Surface-Based Analysis: I. Segmentation and Surface Reconstruction.” *NeuroImage* 9 (2): 179–94. https://doi.org/10.1006/nimg.1998.0395.

Esteban, Oscar, Ross Blair, Christopher J. Markiewicz, Shoshana L. Berleant, Craig Moodie, Feilong Ma, Ayse Ilkay Isik, et al. 2018. “FMRIPrep.” *Software*. Zenodo. https://doi.org/10.5281/zenodo.852659.

Esteban, Oscar, Christopher Markiewicz, Ross W Blair, Craig Moodie, Ayse Ilkay Isik, Asier Erramuzpe Aliaga, James Kent, et al. 2018. “fMRIPrep: A Robust Preprocessing Pipeline for Functional MRI.” *Nature Methods*. https://doi.org/10.1038/s41592-018-0235-4.

Evans, AC, AL Janke, DL Collins, and S Baillet. 2012. “Brain Templates and Atlases.” *NeuroImage* 62 (2): 911–22. https://doi.org/10.1016/j.neuroimage.2012.01.024.

Fonov, VS, AC Evans, RC McKinstry, CR Almli, and DL Collins. 2009. “Unbiased Nonlinear Average Age-Appropriate Brain Templates from Birth to Adulthood.” *NeuroImage* 47, Supplement 1: S102. https://doi.org/10.1016/S1053-8119(09)70884-5.

Glasser, Matthew F., Stamatios N. Sotiropoulos, J. Anthony Wilson, Timothy S. Coalson, Bruce Fischl, Jesper L. Andersson, Junqian Xu, et al. 2013. “The Minimal Preprocessing Pipelines for the Human Connectome Project.” *NeuroImage*, Mapping the connectome, 80: 105–24. https://doi.org/10.1016/j.neuroimage.2013.04.127.

Gorgolewski, K., C. D. Burns, C. Madison, D. Clark, Y. O. Halchenko, M. L. Waskom, and S. Ghosh. 2011. “Nipype: A Flexible, Lightweight and Extensible Neuroimaging Data Processing Framework in Python.” *Frontiers in Neuroinformatics* 5: 13. https://doi.org/10.3389/fninf.2011.00013.

Gorgolewski, Krzysztof J., Oscar Esteban, Christopher J. Markiewicz, Erik Ziegler, David Gage Ellis, Michael Philipp Notter, Dorota Jarecka, et al. 2018. “Nipype.” *Software*. Zenodo. https://doi.org/10.5281/zenodo.596855.

Greve, Douglas N, and Bruce Fischl. 2009. “Accurate and Robust Brain Image Alignment Using Boundary-Based Registration.” *NeuroImage* 48 (1): 63–72. https://doi.org/10.1016/j.neuroimage.2009.06.060.

Huntenburg, Julia M. 2014. “Evaluating Nonlinear Coregistration of BOLD EPI and T1w Images.” Master’s Thesis, Berlin: Freie Universität. http://hdl.handle.net/11858/00-001M-0000-002B-1CB5-A.

Jenkinson, Mark, Peter Bannister, Michael Brady, and Stephen Smith. 2002. “Improved Optimization for the Robust and Accurate Linear Registration and Motion Correction of Brain Images.” *NeuroImage* 17 (2): 825–41. https://doi.org/10.1006/nimg.2002.1132.

Klein, Arno, Satrajit S. Ghosh, Forrest S. Bao, Joachim Giard, Yrjö Häme, Eliezer Stavsky, Noah Lee, et al. 2017. “Mindboggling Morphometry of Human Brains.” *PLOS Computational Biology* 13 (2): e1005350. https://doi.org/10.1371/journal.pcbi.1005350.

Lanczos, C. 1964. “Evaluation of Noisy Data.” *Journal of the Society for Industrial and Applied Mathematics Series B Numerical Analysis* 1 (1): 76–85. https://doi.org/10.1137/0701007.

Power, Jonathan D., Anish Mitra, Timothy O. Laumann, Abraham Z. Snyder, Bradley L. Schlaggar, and Steven E. Petersen. 2014. “Methods to Detect, Characterize, and Remove Motion Artifact in Resting State fMRI.” *NeuroImage* 84 (Supplement C): 320–41. https://doi.org/10.1016/j.neuroimage.2013.08.048.

Reuter, Martin, Herminia Diana Rosas, and Bruce Fischl. 2010. “Highly Accurate Inverse Consistent Registration: A Robust Approach.” *NeuroImage* 53 (4): 1181–96. https://doi.org/10.1016/j.neuroimage.2010.07.020.

Satterthwaite, Theodore D., Mark A. Elliott, Raphael T. Gerraty, Kosha Ruparel, James Loughead, Monica E. Calkins, Simon B. Eickhoff, et al. 2013. “An improved framework for confound regression and filtering for control of motion artifact in the preprocessing of resting-state functional connectivity data.” *NeuroImage* 64 (1): 240–56. https://doi.org/10.1016/j.neuroimage.2012.08.052.

Treiber, Jeffrey Mark, Nathan S. White, Tyler Christian Steed, Hauke Bartsch, Dominic Holland, Nikdokht Farid, Carrie R. McDonald, Bob S. Carter, Anders Martin Dale, and Clark C. Chen. 2016. “Characterization and Correction of Geometric Distortions in 814 Diffusion Weighted Images.” *PLOS ONE* 11 (3): e0152472. https://doi.org/10.1371/journal.pone.0152472.

Tustison, N. J., B. B. Avants, P. A. Cook, Y. Zheng, A. Egan, P. A. Yushkevich, and J. C. Gee. 2010. “N4ITK: Improved N3 Bias Correction.” *IEEE Transactions on Medical Imaging* 29 (6): 1310–20. https://doi.org/10.1109/TMI.2010.2046908.

Wang, Sijia, Daniel J. Peterson, J. C. Gatenby, Wenbin Li, Thomas J. Grabowski, and Tara M. Madhyastha. 2017. “Evaluation of Field Map and Nonlinear Registration Methods for Correction of Susceptibility Artifacts in Diffusion MRI.” *Frontiers in Neuroinformatics* 11. https://doi.org/10.3389/fninf.2017.00017.

Zhang, Y., M. Brady, and S. Smith. 2001. “Segmentation of Brain MR Images Through a Hidden Markov Random Field Model and the Expectation-Maximization Algorithm.” *IEEE Transactions on Medical Imaging* 20 (1): 45–57. https://doi.org/10.1109/42.906424
